# Defoliation estimation of forest trees from ground-level images

**DOI:** 10.1101/441733

**Authors:** Ursula Kälin, Nico Lang, Christian Hug, Arthur Gessler, Jan Dirk Wegner

## Abstract

In this paper, we propose to estimate tree defoliation from ground-level RGB photos with convolutional neural networks (CNN). Tree defoliation is usually assessed with field campaigns, where experts estimate multiple tree health indicators per sample site. Campaigns span entire countries to come up with a holistic, nation-wide picture of forest health. Surveys are very laborious, expensive, time-consuming and need a large number of experts. We aim at making the monitoring process more efficient by casting tree defoliation estimation as an image interpretation problem. What makes this task challenging is strong variance in lighting, viewpoint, scale, tree species, and defoliation types. Instead of accounting for each factor separately through explicit modelling, we learn a joint distribution directly from a large set of annotated training images following the end-to-end learning paradigm of deep learning. The proposed workflow works as follows: (i) Human experts visit individual trees in forests distributed all over Switzerland, (ii) acquire one photo per tree with an off-the-shelf, hand-held RGB camera and (iii) assign a defoliation value. The CNN approach is (iv) trained on a subset of the images with expert defoliation assessments and (v) tested on a hold-out part to check predicted values against ground truth. We evaluate our supervised method on three data sets with different level of difficulty acquired in Swiss forests and achieve an average mean absolute error (avgMAE) of 7.6% for the joint data set after cross-validation. Comparison to a group of human experts on one of the data sets shows that our CNN approach performs only 0.9 percent points worse. We show that tree defoliation estimation from ground-level RGB images with a CNN works well and achieves performance close to human experts.

## 1. Introduction

Forests are of vital importance to human life and the environment in general because they provide a vast array of ecosystem services, from global to local scales, store large amounts of carbon, and by continued sequestration they strongly contribute to the terrestrial carbon sink (Pan et al., 2011). In addition, forests modulate local to global climate via their albedo, surface roughness and their influence on the water cycle (Bonan, 2008). Moreover, they provide timber as construction material and fibers for bioenergy, paper production and the chemical industry (Richardson et al., 2006), and are also biodiversity hotspots (Foley et al., 2005). However, forest functions and their provided services are threatened by various factors such as land use change, air pollutants and climate change (Millar and Stephenson, 2015). Since the 1980s much effort has gone into large-scale forest monitoring in Europe to understand the impact of environmental pollution and a changing climate on tree health (Lorenz, 1995). Defoliation is a main indicator of tree health and is usually assessed by experts that survey tree stands visually in the field at selected sites across the whole country (e.g., a 16 × 16 *km* grid in Switzerland (Dobbertin and Brang, 2001)). A lot of research has been done to automate parts of this tedious, time-consuming, and costly monitoring process. Many works aim at detecting trees and classifying them into their species (Larsen et al., 2011; Kaartinen et al., 2012; Hovi et al., 2016; Blomley et al., 2017). In this regard, the primary data source is remote sensing like multi-spectral aerial (Leckie et al., 2005; Waser et al., 2011) or satellite images (Pu and Landry, 2012), hyper-spectral data (Clark et al., 2005; Roth et al., 2015), dense (full-waveform) LiDAR point clouds (Brandtberg, 2007; Yao et al., 2012b; Hovi et al., 2016; Blomley et al., 2017), or a combination of LiDAR and multispectral images (Heikkinen et al., 2011; Korpela et al., 2011; Heinzel and Koch, 2012). Unmanned aerial vehicles (UAVs) provide a more flexible way of analysing forests. They can acquire data with very high temporal and spatial resolutions for a pre-defined site of interest. Moreover, UAVs can be equipped with different sensors like multi-spectral cameras (Dash et al., 2017) or hyperspectral and LiDAR sensors (Sankay et al., 2017).

Another option, which we follow in our work presented in this paper, is analysis of photos acquired on the ground (Du et al., 2007; Kumar et al., 2012; Mouine et al., 2013; Goëau et al., 2013, 2014; Wegner et al., 2016; Branson et al., 2018). Ground-level images have higher spatial resolution than remote sensing data and provide a horizontal view on the entire tree comparable to the view of the expert assessing the tree visually in the field (Solberg and Strand, 1999). They can be acquired in a more flexible way and do not require expensive, dedicated flight campaigns. This also allows to enthuse citizen scientists for helping with the monitoring effort, for example, through mobile phone apps like *Pl@ntNet* (Goëau et al., 2013, 2014), *Leafsnap* (Kumar et al., 2012), or iNaturalist (http://www.inaturalist.org).

A downside of ground-level photos acquired with off-the-shelf consumer grade cameras is the absence of an infrared channel, which contributes most evidence for tree health assessment from aerial or satellite imagery (e.g., (Huete et al., 1997)). Here, we investigate whether we can estimate defoliation of forest trees from ground-level images based solely on RGB image texture. We use convolutional neural networks (CNNs) as our core component, which have seen huge success for a wide range of applications since the comeback of deep learning (Krizhevsky et al., 2012). In contrast to traditional image interpretation that builds on hand-crafted (texture) features, CNNs learn a rich set of discriminative features directly from the data. CNNs have further properties like translation invariance and robustness to (slight) changes of viewpoint and scale that are very useful in our case. We adapt a ResNet (He et al., 2016) CNN architecture to the task and evaluate our method on three different image data sets. All images show forest trees in Switzerland acquired with hand-held photo cameras on the ground. Photos were captured during field surveys and come with expert labels for defoliation. Our results indicate that automated image interpretation with CNNs performs very well for estimating tree defoliation, close to human performance. We view the work presented here as a first step towards an automated tool that can estimate tree health in order to (i) make professional field campaigns less costly, (ii) facilitate quicker collection of much larger data sets, (iii) and enable contributions by citizen scientists via building our system into an app.

## 2. Related work

*Tree stress estimation* is usually approached either by expert assessments in situ or by analysis of overhead images that include infra-red and near-infrared channels. A good overview of in situ tree monitoring is given by Eichhorn et al. and Morgenroth and Östberg (2017) whereas a comprehensive review of forest health assessment methods with remote sensing is given by Lausch et al. (2016); Lausch et al.. In situ tree monitoring in forests often comprises the visual assessment of defoliation and the resulting increase in crown transparency together with estimates of leaf discolouration. In remote sensing, usually multi-spectral imagery, hyperspectral data or airborne laser scanning are used to estimate health indicators at the level of individual trees. For example, Polewski et al. (2015b) introduce a hierarchical workflow that detects single standing dead trees from aerial color infrared imagery with shape and intensity priors. A Gaussian Mixture Model clusters handcrafted features into categories dead trees, living trees, and shadows. Dead tree regions are refined with an interative level set segmentation method. The same authors propose another multi-step approach for the same task but combine full waveform LiDAR point clouds and color infrared aerial images (Polewski et al., 2015a). 3D point clouds are used to segment individual trees and to generate tree boundary polygons in the images. After projecting LiDAR points into individual tree polygons, they extract hand-crafted features and employ a logistic regression to classify dead trees and living trees. Eitel et al. (2011) use red-edge information of the RapidEye satellite constellation to detect stress of forest trees from space. They compare different indices like the Normalized Difference Vegetation Index (NDVI) and Green NDVI against the Normalized Difference Red-Edge Index (NDRE) and find the latter to perform best. Very high-resolution drone images provide additional means to monitor forest health via spectral indices (Dash et al., 2017). Another widely used method to estimate tree stress is airborne laser scanning in combination with a supervised classification. Shendryk et al. (2016) combine full-waveform airborne laser scans with hyperspectral data and train a Random Forest classifier to estimate dieback and transparency. Mak and Hu (2014) determine the health status of ash trees with ground-based mobile laser scanning as a function of the point density. Lin et al. (2014) apply hyperspectral imagery to the task. They identify if tree leafs change color and humidity depending on their health status and season.

*Tree defoliation estimation* is a major component of tree stress estimation and especially the laser scanning methods mentioned previously are primarily based on estimating defoliation (Yao et al., 2012a; Lin et al., 2014; Shendryk et al., 2016). A wide variety of research applies remote sensing to defoliation estimation of forest trees (Kantola et al., 2010; Mozgeris and Augustaitis, 2013; Marx and Kleinschmit, 2017; Hawrylo et al., 2018). Kantola et al. (2010) combine data from airborne laser scanning and aerial images, Mozgeris and Augustaitis (2013) use only aerial images, Marx and Kleinschmit (2017) analyse RapidEye satellite imagery, and Hawrylo et al. (2018) rely on Sentinel-2 satellite images. All works have in common that they propose a traditional supervised classification approach that first extracts a small set of hand-crafted features, usually with spectral indices as their main component, which is then classified with Random Forests, for example.

Estimating tree defoliation from ground-level images has a long history, too, starting with early works like (Lee et al., 1983). Further works of Mizoue (2002) and Dobbertin et al. (2004, 2005) design multi-step image processing workflows that sequentially detect tree regions in the images, extract features, and finally classify often in a semi-supervised way. All methods have in common that they do not apply to the raw image, but need exact delineation of the tree beforehand (Borianne et al., 2017), which is error-prone and often involves user interaction.

In this work, we present a completely automated approach that learns tree defoliation directly from the original images without any previous tree silhouette extraction. To the best of our knowledge, our work is also the first to propose convolutional neural networks for tree defoliation estimation from images. Moreover, most related works test their methods on very small data sets since collection of ground truth stress data is very tedious. For example, the data set of Shendryk et al. (2016) consists of only 54 individual trees of a single species (eucalypt) and Eitel et al. (2011) validate their approach on a single test site with only conifers. In contrast, we train and validate our method on three data sets with over 2000 images of a large variety of different species, shapes, and complex scenarios of different, dense forests. Additionally, we solely rely on standard RGB imagery acquired with hand-held cameras on the ground. We compensate the lack of pixel-wise spectral information or height values with the power of deep CNNs that can learn discriminative texture patterns directly from large amounts of training data. It turns out that CNNs with their very high modelling capacity can learn very complex multi-variate distributions across different tree species, shapes, lighting conditions, varying scale and viewpoint to estimate defoliation with an accuracy similar to human experts.

## 3. Method for Defoliation estimation

Estimating tree defoliation from ground-level RGB photos acquired with off-the-shelf consumer grade cameras is challenging. Image interpretation has to completely rely on texture, viewpoint and lighting variations have to be compensated, and it has to work across a wide range of tree species. Traditional methods that rely on hand-crafted features (Lee et al., 1983; Mizoue, 2002; Dobbertin et al., 2004, 2005; Borianne et al., 2017) have to carefully compensate for all disturbing effects with their model design. In contrast, CNNs learn very high-dimensional multi-variate distributions over species and defoliation directly. Viewpoint changes are compensated via built-in translation invariance. A rich set of learned filters in combination with activation functions (rectified linear units in our case, abbreviated with ReLU) result in discriminative, non-linear texture representations. We therefore choose CNNs for tree defoliation estimation because they learn features and the classification model jointly end-to-end for a specific task and training data set. Circumnavigating manual feature design by learning the most discriminative features directly from the given data is a major reason for success in comparison to more traditional methods. Our method works as follows: (i) Human experts visit individual trees in forests distributed all over Switzerland, (ii) acquire one photo per tree with an off-the-shelf, hand-held RGB camera and (iii) assign a defoliation value. (iv) All images are collected and squeezed into square patches to make them amenable to the CNN (an adaption of the ResNet architecture (He et al., 2016)) and (v) data augmentation is done by flipping images etc. Our CNN approach is (vi) trained on a subset of the images with expert defoliation assessments and (vii) tested on a hold-out part to check predicted values against ground truth. Note that this procedure is meant for training and validating our approach with reference data collected by professional arborists capable of assessing tree defoliation manually in the field. Once trained and validated, we aim at using the trained model for an app that can be used by any citizen scientist to take a photo of a tree in a Swiss forest, where tree defoliation prediction is then done automatically by our model.

### 3.1. Data pre-processing

An important step before training is data pre-processing, which involves cropping or down-sampling (images of our data set have been acquired with multiple different cameras and images therefore have different resolutions), normalization of the radiometric distributions, and data augmentation. In a first step, we transform all images into square patches of size 256 × 256 *pixels* in order to achieve a single input image size across all samples. Since all our images are larger and of rectangular shape, squeezing them involves anisotropic downsampling (Fig. 1). Relatively small patches save memory and help staying within the memory budget of our GPU, but still allow for sufficiently high batch sizes (32 in our case). The technical term *batch* refers to a small sub-sample of images from the training data set used to compute the gradients and the loss function during backpropagation. Higher numbers of images per batch allow for higher learning rates, which in turn accelerates training. In order to ensure sufficient detail in the down-sampled images for training tree defoliation, we ran preliminary tests with higher image resolution of 512 × 512 *pixels*. However, we did not observe significant performance gains but training time increased significantly. We thus decided to keep an image size of 256 × 256 *pixels* for all experiments. A second step for pre-processing is the normalization of the image channels to achieve the same radiometric distribution across all input images. We zero-center (subtraction of the mean) and scale (division by the standard deviation) separately for all three channels. Note that mean and standard deviation are computed only on the training set and applied to both, train and test set. The final pre-processing step is data augmentation, which aims at artificially increasing the (labeled) training data set by applying transformations to the existing images. This simulation of additional training samples often helps generalizing the model because the distribution over the training data set is closer to the true distribution. However, this is only valid if we use transformations for augmentation that would naturally occur in the data (but are underrepresented). In our case, we mirror each image along the vertical axis, zoom slightly and add a bit of rotation. This makes the trained model more robust against viewpoint changes, scaling effects, and small tilt variations of either camera or tree.

**Figure 1:**
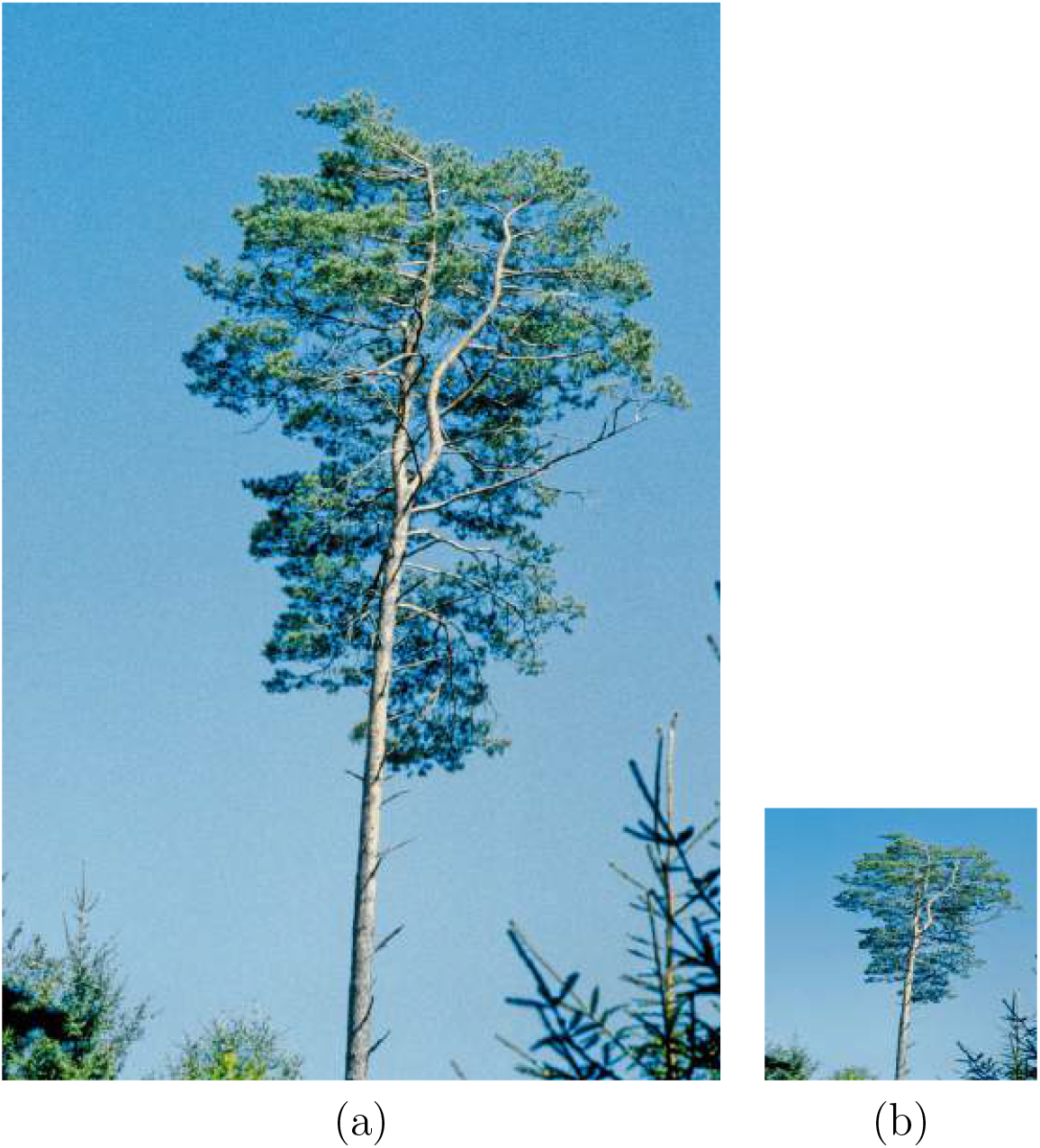
Example from the *Urmeter* data set of (a) the original image (675 × 1013 *pixels*) and (b) image patch (256 × 256 *pixels*) used as input to the CNN after anisotropic down-scaling.

### 3.2. Architecture of the CNN

We train a standard ResNet architecture (He et al., 2016) for regression to estimate tree defoliation for all experiments. ResNet is current state-of-the-art in image classification and in the following we briefly remind the reader of its main technical advantages over alternative architectures like VGG (Simonyan and Zisserman, 2015), for example. The ResNet architecture allows to generate deeper networks that behave well during the training processes. Usually, the construction of deeper networks results in dramatic increase in number of weights as well as exploding or vanishing gradients during the learning process. ResNet reduces this problem significantly with a specific design of convolutional building blocks, where each block contains two branches. One is propagated through the layers as usual, whereas the other branch is used as an identity mapping that is added again at the end of each building block (see the original paper of He et al. (2016) for an in-depth explanation and illustrations). Thus, the network learns a residual function, which behaves more stable during the training process. In this work, we use a ResNet50 architecture, which consists of 16 convolutional blocks in total. Our architecture is shown in Tab. 1. Convolutional building blocks always consist of a series of image convolutions with filters of relatively small kernel size (maximum 7 × 7 *pixels* for *conv1*). Naturally, the filter number per convolutional layer is identical to the number of output activation maps. Each building block can be repeated multiple times in a row (e.g., six times *conv4*). Max pooling spatially down-scales the output of *conv1* by replacing the nine values in the 3 × 3 kernel with their maximum one. Recall that the average pooling operation before *fc1* works like max pooling, but uses the maximum value instead of the average across the kernel. The final, fully-connected layer *fc2* generates a 1-dimensional vector and results in a single output, i.e. the defoliation estimate for the input image patch.

**Table 1:**
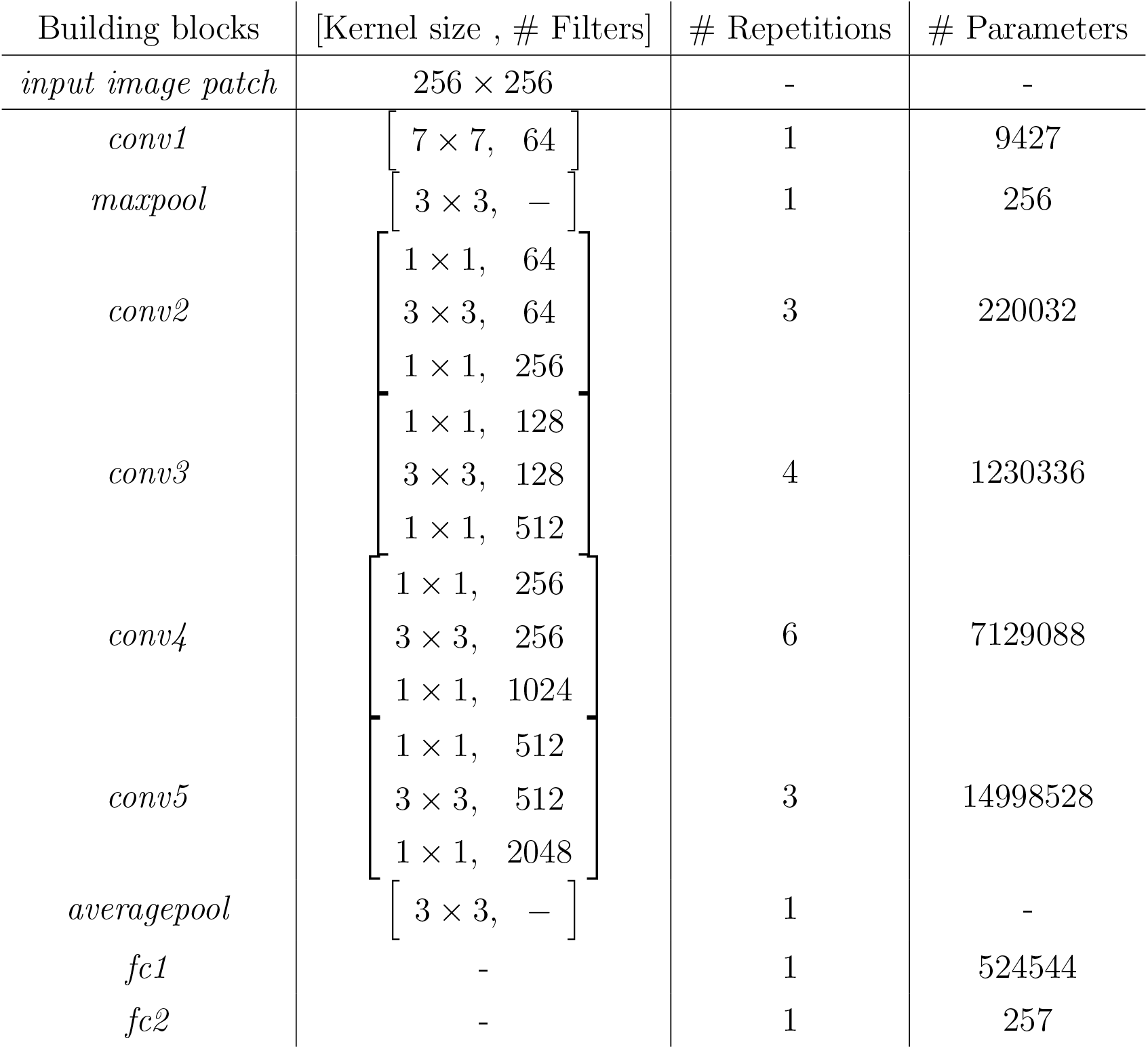
ResNet architecture (He et al., 2016) as used for all our experiments with 24112513 parameters in total. # Repetitions refers to the number each building block (in square brackets) is executed sequentially (e.g., *conv2* runs three times in a row).

| Building blocks | [Kernel size , # Filters] | # Repetitions | # Parameters |
| --- | --- | --- | --- |
| *input image patch* | $256 \times 256$ | - | - |
| *conv1* | $\begin{bmatrix} 7 \times 7, & 64 \end{bmatrix}$ | 1 | 9427 |
| *maxpool* | $\begin{bmatrix} 3 \times 3, & - \end{bmatrix}$ | 1 | 256 |
| *conv2* | $\begin{bmatrix} 1 \times 1, & 64 \\ 3 \times 3, & 64 \\ 1 \times 1, & 256 \end{bmatrix}$ | 3 | 220032 |
| *conv3* | $\begin{bmatrix} 1 \times 1, & 128 \\ 3 \times 3, & 128 \\ 1 \times 1, & 512 \end{bmatrix}$ | 4 | 1230336 |
| *conv4* | $\begin{bmatrix} 1 \times 1, & 256 \\ 3 \times 3, & 256 \\ 1 \times 1, & 1024 \end{bmatrix}$ | 6 | 7129088 |
| *conv5* | $\begin{bmatrix} 1 \times 1, & 512 \\ 3 \times 3, & 512 \\ 1 \times 1, & 2048 \end{bmatrix}$ | 3 | 14998528 |
| *averagepool* | $\begin{bmatrix} 3 \times 3, & - \end{bmatrix}$ | 1 | - |
| *fc1* | - | 1 | 524544 |
| *fc2* | - | 1 | 257 |

Defoliation is estimated as a continuous value and we thus have to solve a regression task. Two rather simple loss functions are usually applied for regression that either minimize the norm of the difference between prediction and ground truth during training (L1-loss) or the squared norm (L2-loss). Note that L1 basically translates to minimizing the mean absolute error whereas L2 minimizes the mean squared error. We tried both L1 and L2 but did not find a significant performance gap. We thus opt for the L2-loss for all our experiments, which is the most widely used loss function for regression.

### 3.3. Training the CNN

CNNs are trained via backpropagation that updates network weights along high gradients (steepest decrease of the loss) to minimize the discrepancy between predicted defoliation (computed with a forward pass) and the reference label. A quasi-standard today is mini-batch stochastic gradient descent. Its main idea is to randomly sample a small subset from the training data set (32 images per batch in our case) and doing backpropagation repeatedly, alternating with forward passes. A new mini-batch of training images is sampled for each backpropagation. An important hyper-parameter for tuning the learning process is the learning rate. The learning rate defines how much weight we give to the gradients computed during backpropagation to update the trainable weights. With a reasonably large batch size of 32 images that can be assumed to well represent the distribution of the entire training data set, we can apply higher learning rates resulting in more rapid decrease of the loss. In contrast, smaller batch sizes (e.g., only four images) result in less memory consumption on the GPU. However, this comes at a cost: we cannot trust the subset of images to be a representative sample of the full training data set as much and thus have to set a smaller learning rate. This results in much longer training times. Moreover, a smaller learning rate risks the solver getting stuck in local minima early on during the training process. We thus always aim at fitting a larger batch size into GPU memory because this allows for smoother optimization and it globally accelerates training.

Often, the learning rate is decayed during the course of training. At the beginning, bigger steps are needed to quickly move towards a favorable solution without getting stuck in local minima, whereas small steps towards the end prevent the training procedure from fluctuating around minima. Early optimizers use the same learning rate for adjusting all weights and were hard to steer to achieve a stable training process. Newer implementations of optimizers such as Adagrad (Duchi et al., 2011), RMSProp (Hinton et al., 2012), or Adam (Kingma and Ba, 2014) include complexer strategies that can handle the decay and are capable of tuning the learning rate for each learnable parameter individually. We choose to use Adam as optimizer for all experiments because Kingma and Ba (2014) show that it outperforms RMSProp towards the end of the learning process when gradients become sparser.

## 4. Experiments

Visual inspection of trees is very challenging due to their vast variation in species, shape, height, texture and color. Although species-specific models could potentially predict tree stress more accurately, we lack sufficient training data per species to train a strong model. We thus build a single, species-agnostic model to predict individual tree health from ground-level images. Since deciduous trees lose their leafs in winter, visual health indicators like the dieback, leaf discoloration, and crown transparency are invalid outside the growing season. Therefore, all images were taken between beginning of July and end of August, when also the in situ assessments by the experts are carried out. In the following, we first describe the data sets we use for testing our approach and our evaluation strategy.

### 4.1. Data and Test Site

We evaluate our method on three different data sets acquired during long-term campaigns of the Swiss Federal Research Institute WSL. All data sets have in common that a trained observer took photos of individual trees during a field survey. Photos are mostly well centered on a single tree and captured with an appropriate zoom level. However, due to dense, complex forest scenarios, the tree of interest can be partially occluded, dense forest may appear in its background, and lighting conditions vary largely. We base our experimental evaluation on data sets *Urmeter, Parcours*, and *WSI* that represent different levels of difficulty. A variety of species that are typical for Swiss forests is contained in the data sets (Fig. 2). All three data sets have in common that samples of high defoliation are rare whereas most samples show low defoliation (Fig. 3).

**Figure 2:**
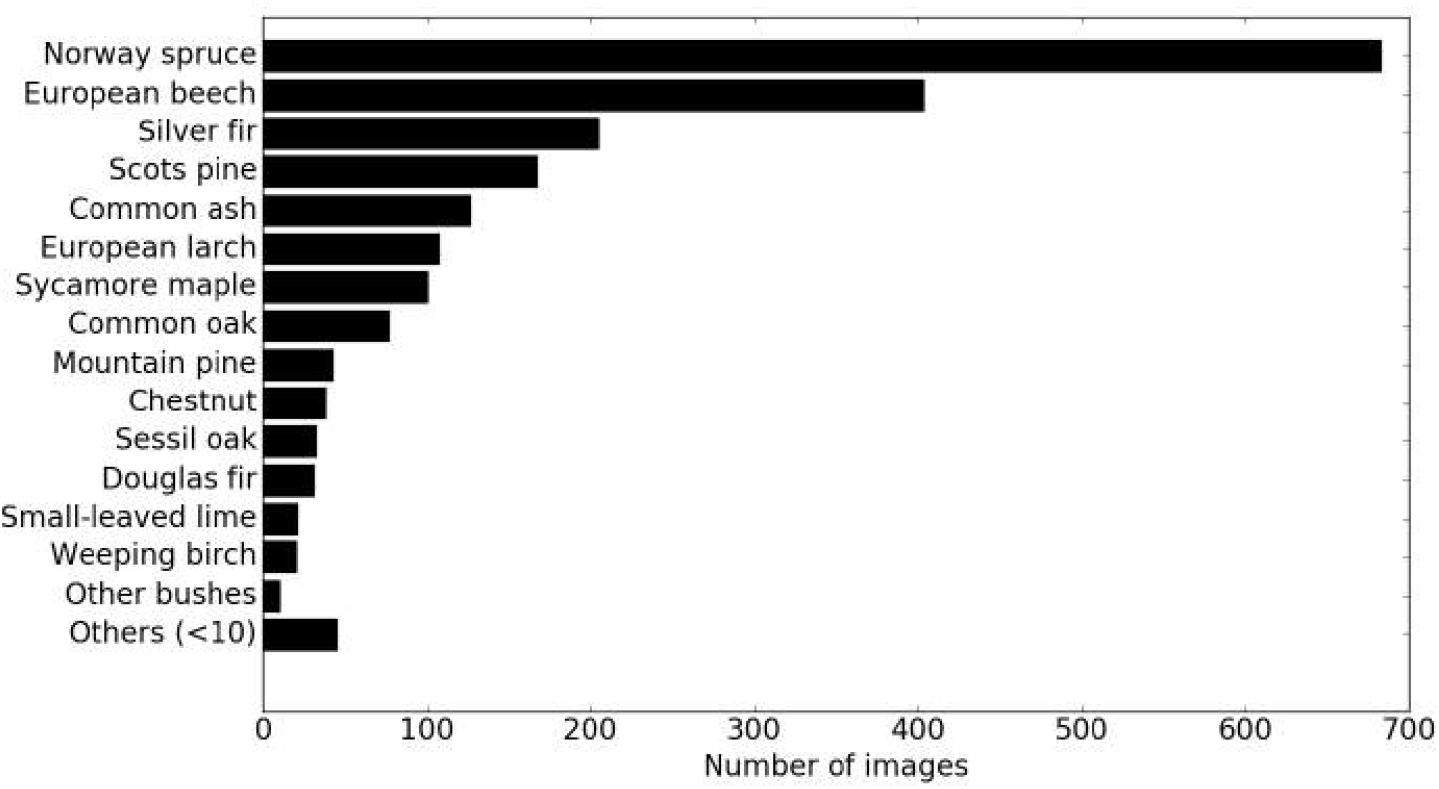
Distribution of tree species over all three data sets *Urmeter, Parcours*, and WSI.

**Figure 3:**
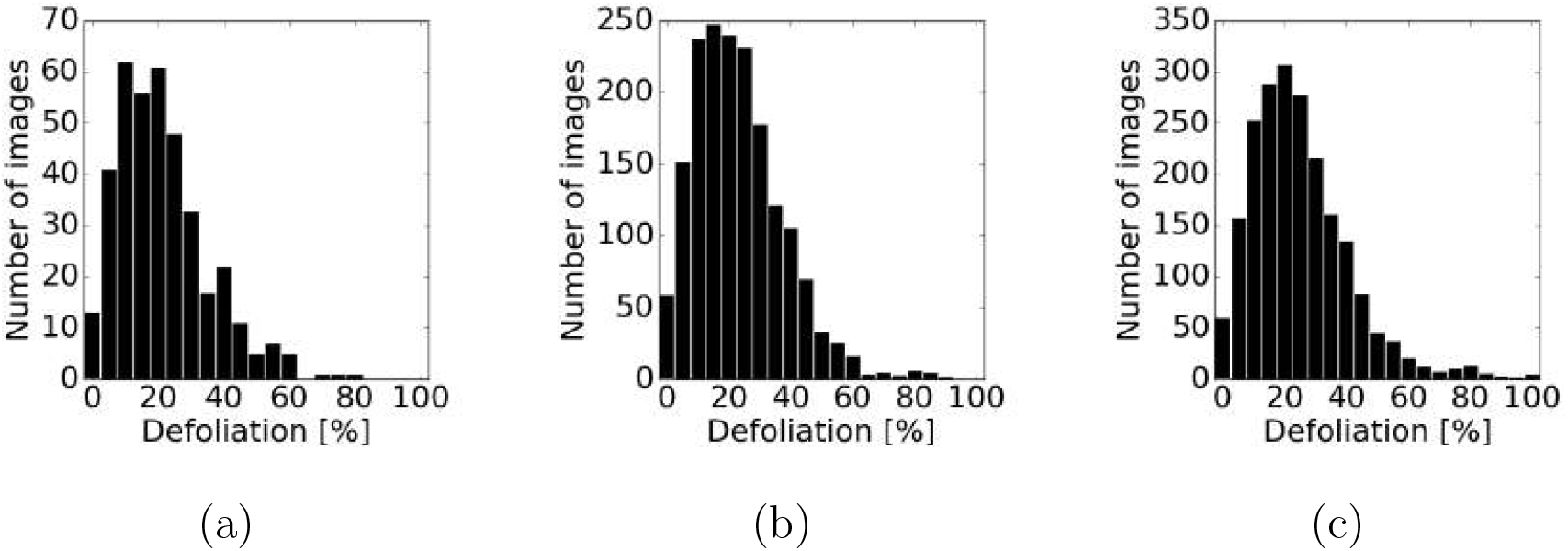
Distribution of samples with respect to defoliation: (a) *Urmeter*, (b) *Parcours*, and (c) WSI.

*Urmeter* consists of 384 images of size 3600×5400 pixels originally recorded as slides from 1987 – 1992 and later scanned. Images contain tree crowns that are standing out of their environment usually with bright sky as background and occluded parts are rare. This set of images was originally constructed as reference image set to teach new observers for field campaigns. Ground truth defoliation labels were estimated by averaging across image-based assessments of six different experts. Defoliation on the images was assessed according to the ICP Forests manual (Eichhorn et al.). Each photo has a label for the defoliation of the target tree in percent (i.e., the relative leaf or needle loss of the given tree compared to a theoretical reference tree with no defoliation). Note that the expert assessment results in values with 5% increments but due to the averaging of assessments of six persons intermediate values occur. This data set serves two different purposes. First, it contains images with easily recognizable, individual trees acquired under favorable lighting conditions (Fig. 4(top row)). Images are relatively easy to interpret and we thus view results with this data set as the best possible or upper bound of what can be expected. Second, the image-based expert assessments serve as a baseline for our automated method. It raises the question of how good a deep learning approach can perform compared to human experts.

**Figure 4:**
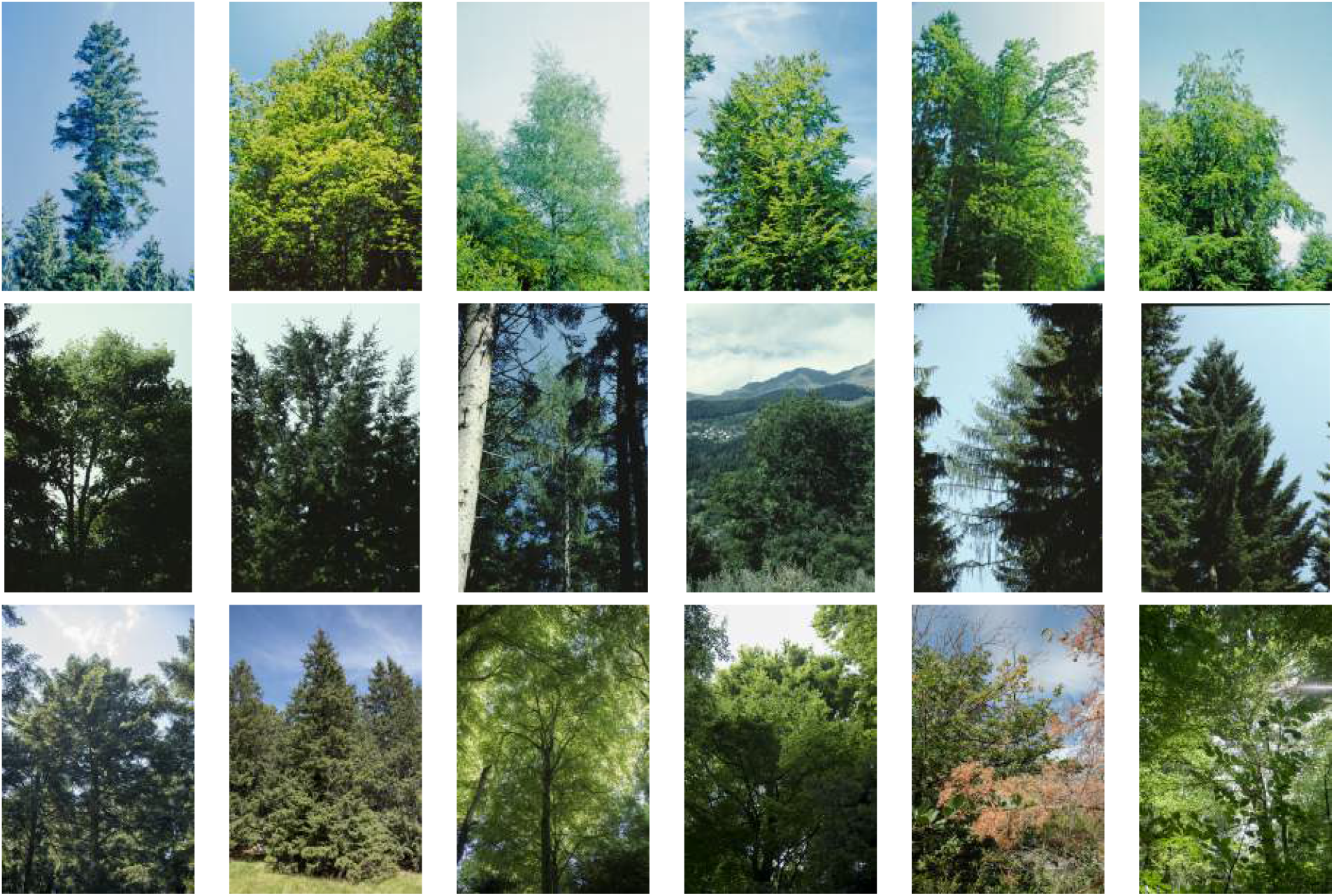
Examples photos of data sets *Urmeter* (top), *Parcours* (center), and *WSI* (bottom).

*Parcours* consists of 1361 images of size 900 × 1370 pixels recorded between 1990 and 1997 originally acquired as slides and later scanned. Images were acquired at ten different locations across Switzerland. Tree defoliation assessments were carried out in the field in contrast to *Urmeter*, where this was done based on the images. Expert assessments were done according to the same protocol as described above. Individual trees are harder to recognize compared to the first data set (*Urmeter*) because they are partially occluded, lighting conditions are less favorable, and denser forest often covers the background (Fig. 4(center row)). It is of intermediate difficulty compared to *Urmeter* (easy) and *WSI* (hard).

*WSI* consists of 363 images of size 3744 × 5616 pixels and 4159 × 6239 pixels recorded with digital cameras in 2017 for the purpose of this study. Photos were taken by experts who estimated defoliation values directly in the field with the same protocol as described above. In contrast to both other data sets, this set of images does not consist of a selection of especially typical or well assessable trees. It rather represents realistic conditions in regular forest plots and includes many images where the tree of interest is highly occluded (Fig. 4(bottom row)). Additionally, trees are mostly photographed with a steeper viewing angle due to less space in front of the tree. Many trees are embedded in dense forest and it is often hard to distinguish individual tree crowns. The radiometric distribution is different (greater variability of brightness and color) with respect to *Urmeter* and *Parcours* because photos were acquired with a modern digital camera instead of scanning analogue slides. Overall, we expect this to be the hardest data set due to steep viewing angle, many occlusions, and dense forest background.

### 4.2. Evaluation strategy

We perform 5-fold cross-validation for our experimental evaluation in order to avoid any train-test split bias. We ensure a roughly equal distribution of defoliation values across all five folds. Total numbers for train and test split per data set are given in Tab. 2. We use the mean absolute error (MAE) to measure the performance of our CNN model on the test data. Because we run 5-fold cross-validation, we provide the average mean absolute error (avgMAE) of the *k* = 5 individual MAE values (Eq. 1). In order to indicate the performance spread across the test data sets, we also provide a standard deviation per experiment.

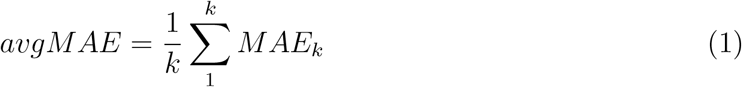

**Table 2:** Number of train and test images per cross-validation fold per data set. Each data set is divided into five mutually exclusive subsets. For each cross-validation fold, four subsets make up the training sample while we apply the trained model to the fifth for testing. Each data subset is once a test set and four times part of the training set.

|  | Train | Test |
| --- | --- | --- |
| *Urmeter* | 307 | 77 |
| *Parcours* | 1088 | 273 |
| *WSI* | 290 | 73 |

Results in Fig. 5 and 6 contain the data points aggregated over all five test splits. We plot prediction (vertical axis) versus ground truth values (horizontal axis) in a 2D-histogram for better visualization of the distribution (instead of a scatter plot). In this visualization, the individual data points are accumulated into 5% bins. The count in each bin is encoded on a grayscale colormap ranging from white (low) to black (high). A regression error characteristic curve (REC curve) (see Fig. 7) is used to evaluate the model accuracy on different error tolerance levels and to compare our CNN approach to human observers.

**Figure 5:**
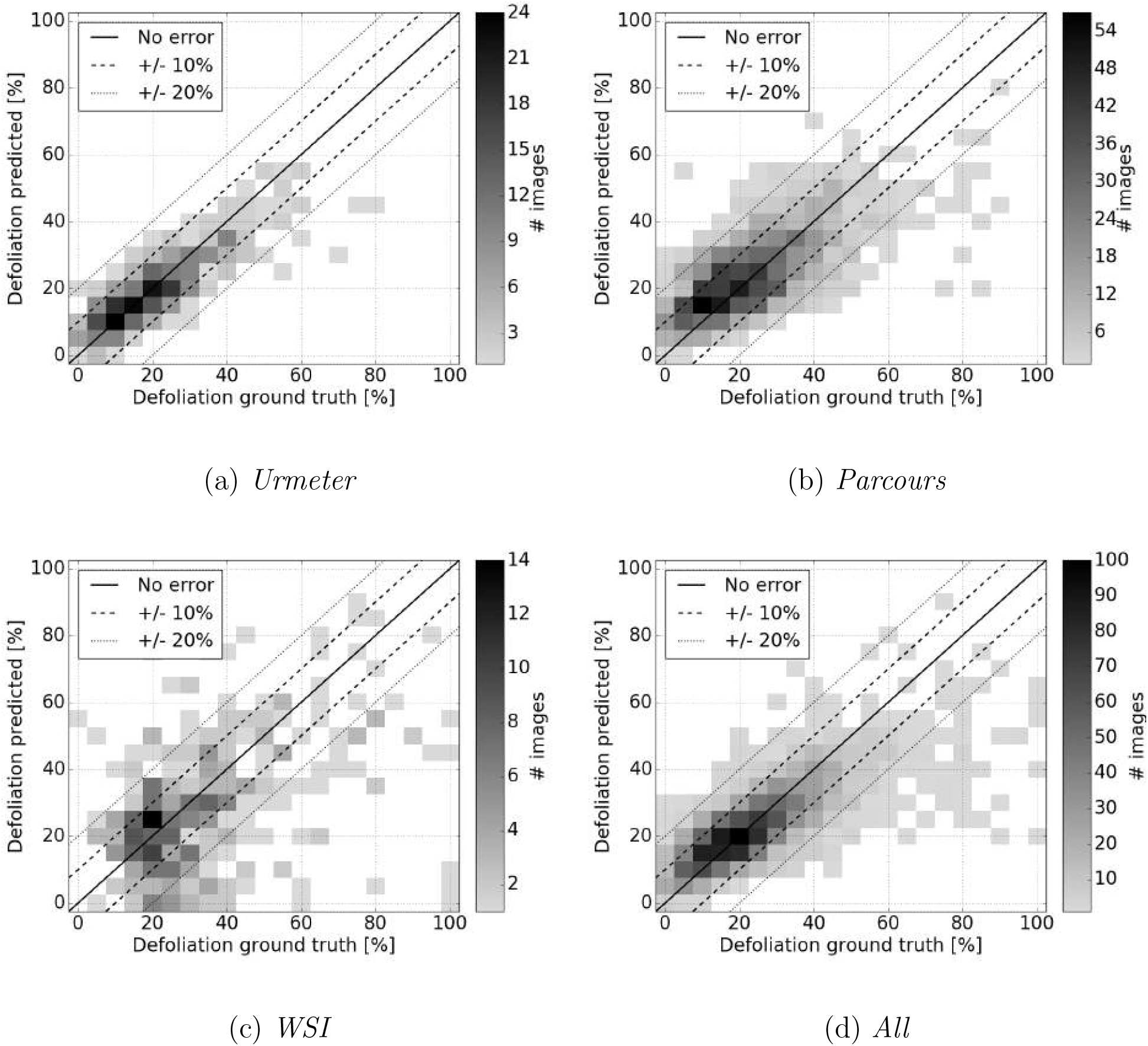
Results of 5-fold cross-validation for training and predicting on *Urmeter* (a), *Parcours* (b), *WSI* (c), and all three data sets jointly (d). Dotted lines indicate error margins in [%] and light grey to black color represents the number of images that fall into an interval (see color bar right of each plot).

We run training per fold for 150 epochs with an initial learning rate of 10^−4^. Preliminary tests showed that using pre-trained weights (on ImageNet) did neither improve results nor reduce training time nor accelerate convergence. The most likely explanation is that our images differ significantly from standard computer vision images in several ways. Our tree images show mostly high-frequency information (leafs, branches) and the loss function aimed at estimating defoliation is focusing on this detailed evidence. Typical computer vision images are dominated by low-frequency evidence, i.e. few objects cover large portions (> 100 *pixels)* of the image. We thus start training from scratch and initialize network weights randomly.

### 4.3. Tree defoliation estimation results

We present results of our experiments with the three data sets in Tab. 3 and Fig. 5. All experiments are run on a NVIDIA GeForce GTX 1080 Ti GPU with 11 GB memory and the approach is implemented using the deep learning library Keras (https://keras.io/). Predicted defoliation is plotted versus the ground truth defoliation. Dotted lines indicate error margins of 10% and 20%, while light grey (few) to black (many) colors encode the number of images that fall into each cell. Prediction results of *Urmeter* (Fig. 5(a)) and *Parcours* (Fig. 5(b)) show a good correlation to the ground truth. Only very few samples of *Urmeter* fall outside the 20% error margin and the majority of images is within the 10% margin (Fig. 3(a)). The average mean absolute error (avgMAE) of *Urmeter* predictions is 5.5%, which is very small.

**Table 3:**
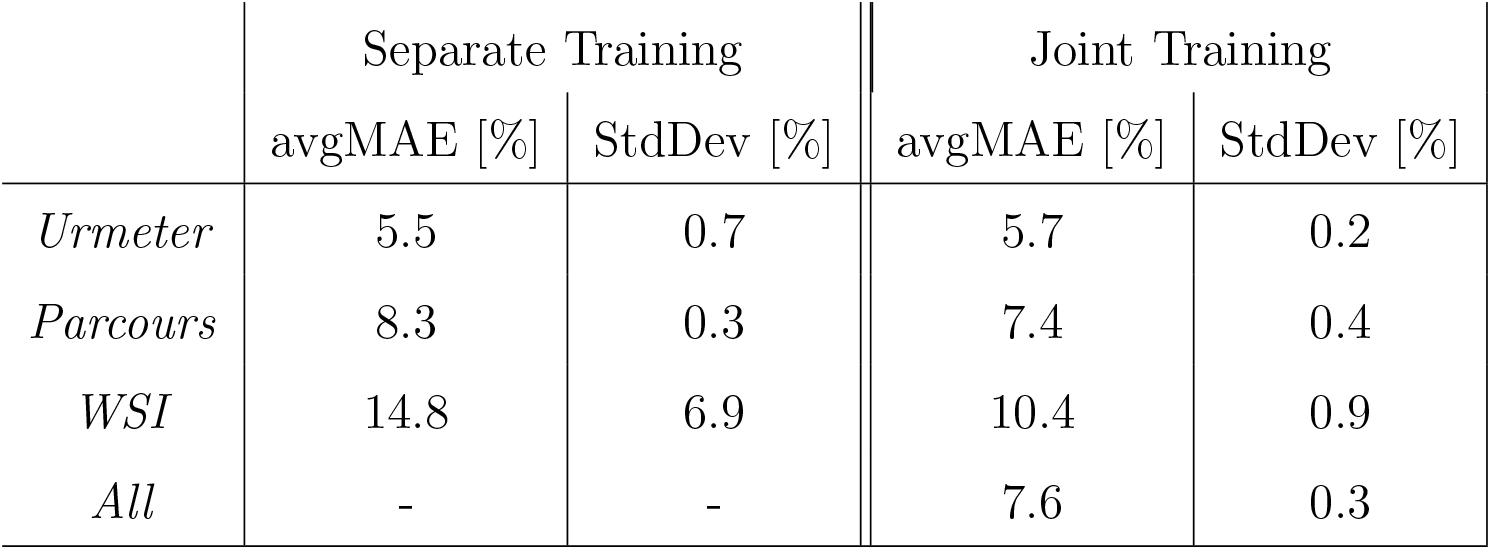
Tree stress estimation results after 5-fold cross-validation for (left) training and predicting on all data sets separately (*Separate Training*) and (right) for training across all data jointly but predicting for the individual data sets (*Joint Training*). Average mean absolute errors (avgMAE) are averaged MAE values across the five test splits per experiment (Eq. 1) with respective standard deviations (StdDev).

|  | Separate Training |  | Joint Training |  |
| --- | --- | --- | --- | --- |
|  | avgMAE [%] | StdDev [%] | avgMAE [%] | StdDev [%] |
| *Urmeter* | 5.5 | 0.7 | 5.7 | 0.2 |
| *Parcours* | 8.3 | 0.3 | 7.4 | 0.4 |
| *WSI* | 14.8 | 6.9 | 10.4 | 0.9 |
| *All* | - | - | 7.6 | 0.3 |

Results for the (harder) *Parcours* data set are slightly worse, but still only few samples fall outside the 20% error margin while most are within 10% deviation from ground truth. The avgMAE of *Parcours* predictions is 8.3% at a very low standard deviation of 0.3%. For images with trees of high defoliation between 80% – 100% there is a tendency to underestimate defoliation (Fig. 5(b)). However, in absolute numbers these gross errors outside the 20% error margin is still low with only 8 images in total. Some of these high defoliation cases are particularly difficult, too, like the example shown in Fig. 8(d) where the actually defoliated part of the tree is occluded in the image by another plant in the foreground. In contrast, results for *WSI* (Fig. 5(c)), the hardest data set with the most complex images, are clearly inferior to both other data sets. Although most images fall into the 20% error margin, a trend that would clearly correlate predictions with ground truth is much weaker. It seems that the amount of training data of *WSI* is too low for learning the multi-variate distributions across all complex scenes. This also leads to a higher mean absolute error of 14.8%. To compensate for this lack of training data we run another experiment where we combine all data sets and train a single model (Fig. 5(d)). Results show that the model can adapt well to the differences across data sets leading to a mean absolute error of 7.6%, which is below the one for *Parcours*.

In order to test whether the larger amount of training data is key to success, we train on the combined data set but predict for each data set separately (Tab. 3 right column *Joint Training*). It turns out that joint training across all three data sets decreases the average mean absolute error from 14.8% to 10.4% for *WSI* and from 8.3% to 7.4% for *Parcours* whereas results of *Urmeter* remain almost unchanged. The standard deviation across all five folds for *WSI* drops significantly from 6.9% to 0.9% while it remains almost unchanged for *Parcours* (0.3% vs. 0.4%). This finding indicates that we can leverage images from other data sets to compensate for lack of data in small data sets. The collection of many more images by WSL in the coming years will further improve results. Jointly training across all data sets does show a slight tendency of underestimating samples with high defoliation. Again, however, in absolute numbers this effect does not play a big role because it amounts to 18 images between 80%–100% defoliation (out of 2108 images in total) that are estimated more than 20 percent points too low (Fig. 5(d)). Four of those images are shown in the top row of Fig. 8, which are particularly challenging cases.

We further compare performance of the deep learning approach to human assessments. Six different experts judged defoliation of trees in the *Urmeter* data set by looking at the images. The average defoliation value across those experts serves as ground truth and individual predictions can be plotted like for all previous experiments (Fig. 6(a)). Results plotted in Fig. 6(a) do not show a big difference compared to the CNN performance in Fig. 5(a). This interpretation is supported by the human avgMAE of 4.6% given in Tab. 4, which is only 0.9 percent points lower than our CNN approach (top left Tab. 3).

**Figure 6:**
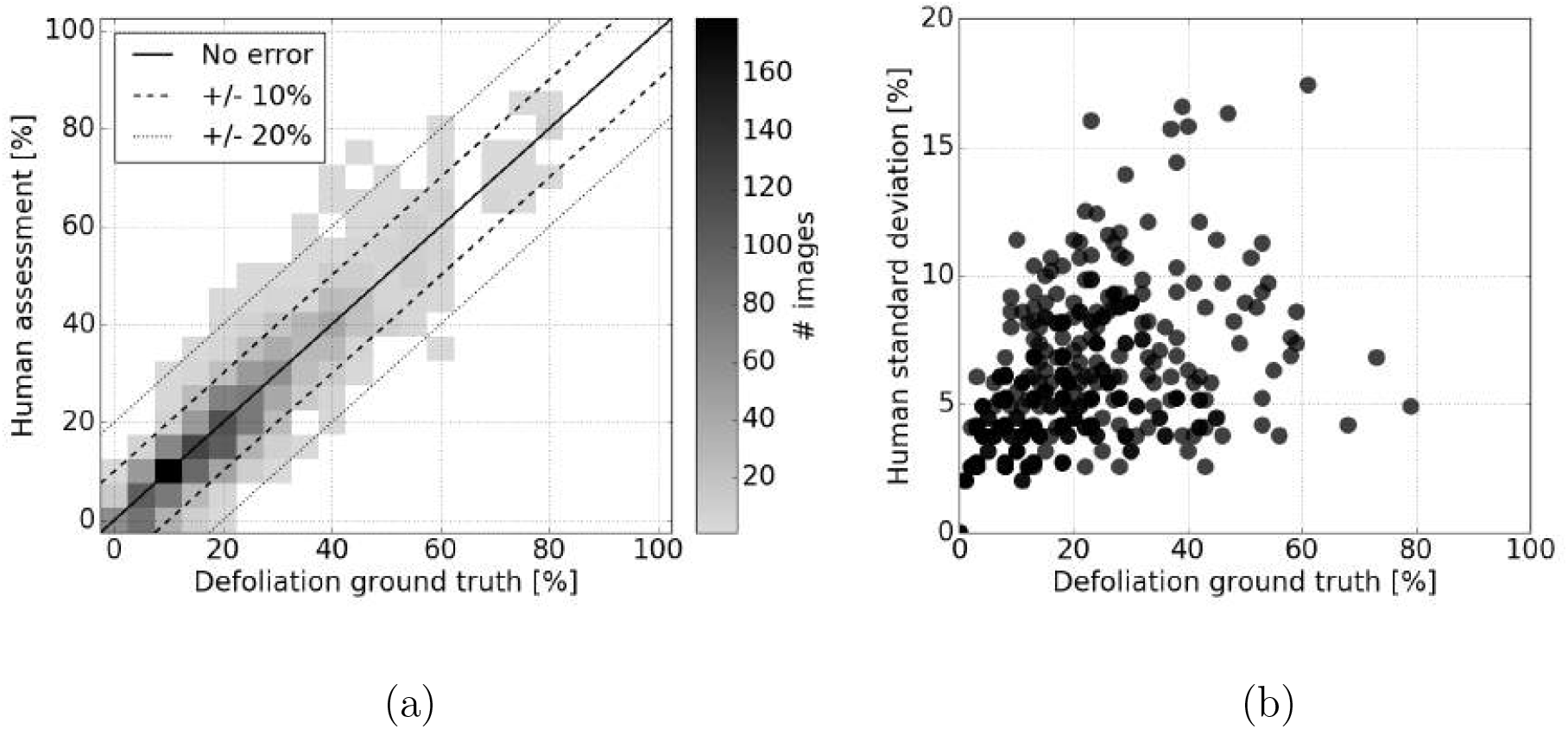
Human expert defoliation assessments on the *Urmeter* data set. (a) Individual human assessments (vertical axis) with respect to ground truth (horizontal axis, average across all six observers), compare to performance of our CNN approach in Fig. 5(a). (b) Standard deviation of individual human assessments (horizontal axis) with respect to the ground truth (vertical axis) for each image in the *Urmeter* data set.

**Table 4:**
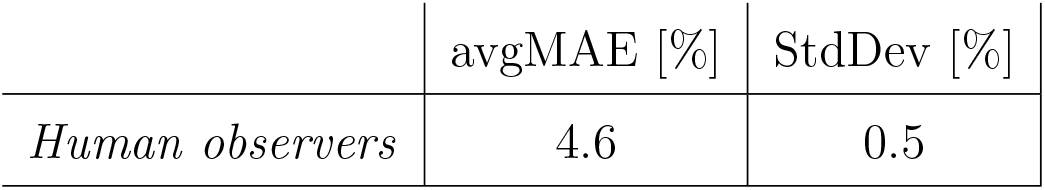
Human performance of 6 individual observers. The average mean absolute error (avgMAE) is calculated with Eq. 1. The standard deviation (StdDev) shows the agreement of the individual observers across the entire data set.

To make the high fluctuation in human expert assessments more obvious, we plot (Fig. 6(b)) the standard deviation across all six human assessments per image with respect to defoliation degrees. This plot indicates that humans are relatively good at assessing very low and very high degrees of defoliation but disagree significantly at intermediate steps (≈ 15% – 60%). We further plot overall accuracy versus error tolerance in Fig. 7, which indicates that the CNN is not far away from human performance. If we allow a 10% error tolerance, our CNN approach achieves only 8 percent points less in overall accuracy than the human experts. The mean absolute error of the human experts allows comparing human defoliation estimation based on images to the ones predicted by the CNN.

**Figure 7:**
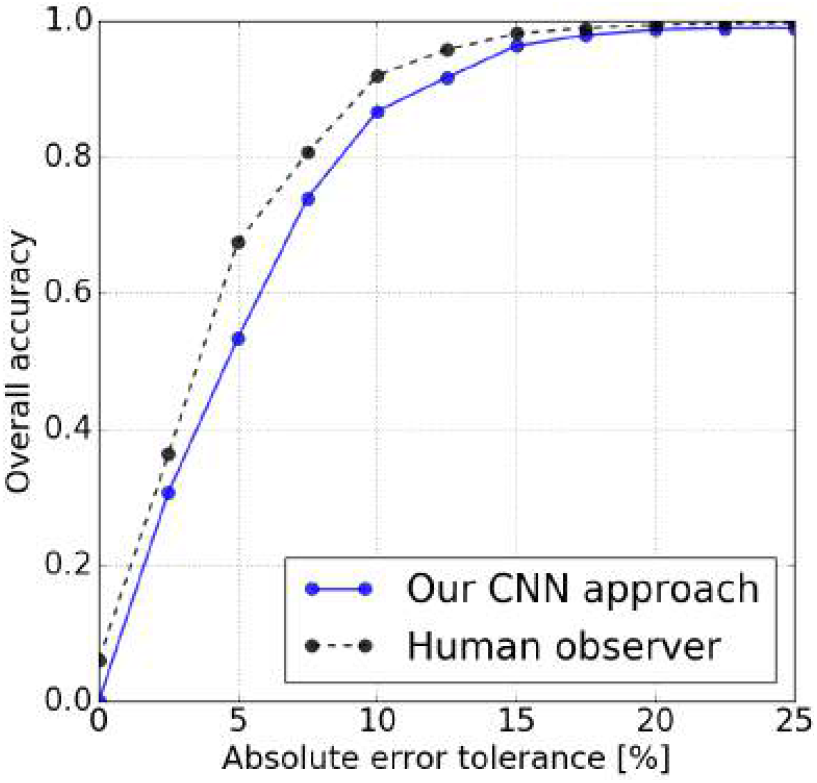
Comparison of human expert defoliation assessments and our CNN approach on the *Urmeter* data set. Accuracies of individual human assessments and our CNN approach with respect to the error tolerance (dotted lines in Fig. 6 (a)).

### 4.4. Discussion of failure cases

In order to better understand the limitations of the automated CNN approach and to discover promising directions for future improvements, we look at the most dominant failure cases as well as examples of images where our method works well. We show a collection of typical images with very high prediction errors in Fig. 8 and good ones in Fig. 9. To support interpretation of results, we also show examples of activation maps in Fig. 10. However, interpreting activation maps directly as evidence for what the CNN learns to look at should be done with care. Recall that there are many activation maps per convolution (e.g., 64 in *conv1* in Tab. 1) and many more if looking at the entire network. The final regression score and result are a non-linear combination of *all* those activation maps for one image, i.e. the true value is in their combination. In this sense, the activation maps we are showing do indicate some cases where, for example, the CNN learns to look at the distribution of branches (Fig. 10(b)), tries to separate tree and sky (Fig. 10(c)), or detects empty space in the canopy (Fig. 10(d)). However, there are many more activation maps per layer that do not allow a direct visual interpretation because they only become valuable in combination with others.

**Figure 8:**
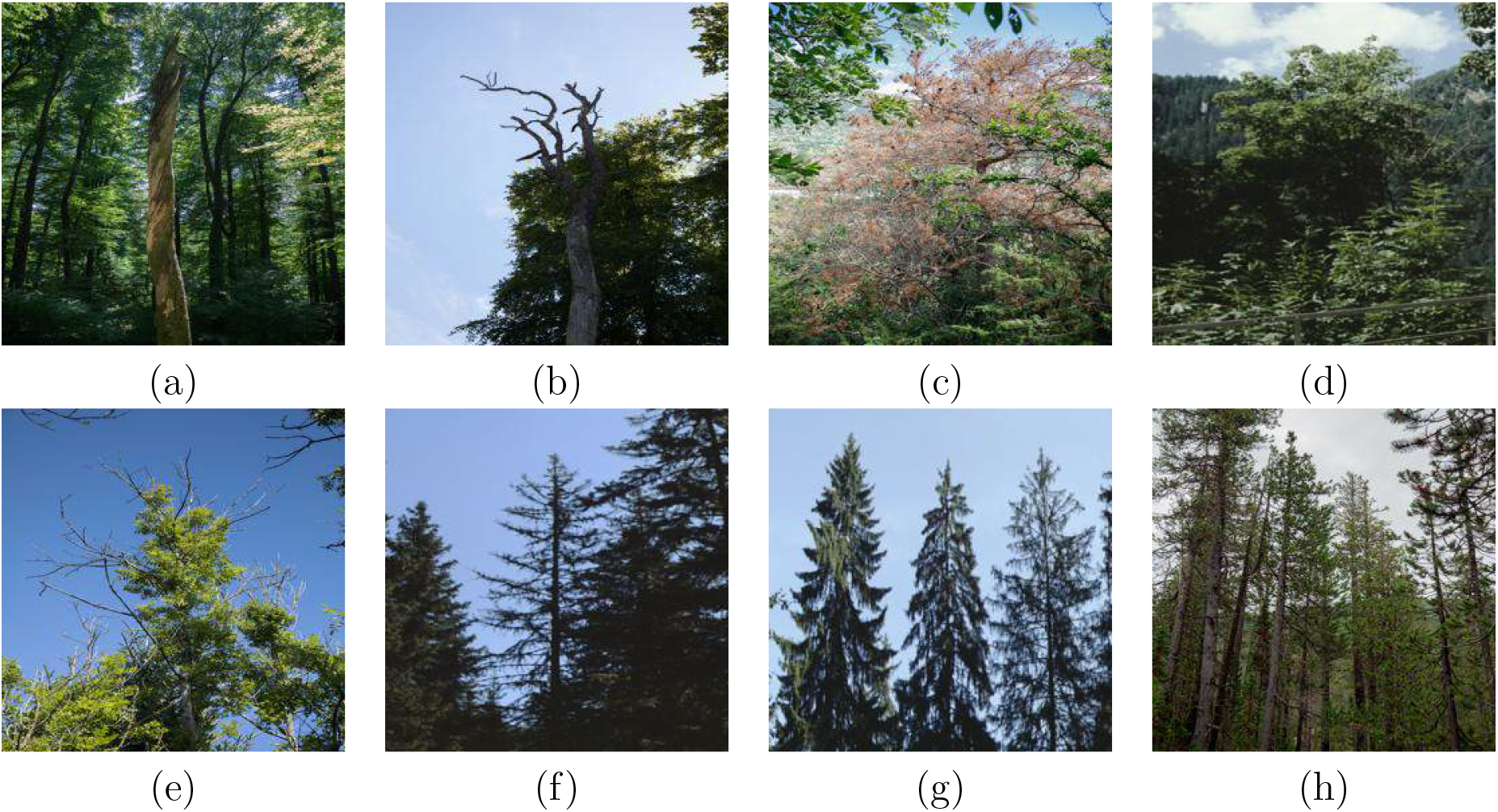
Images with very high prediction errors. The top row shows images where defoliation was predicted too low, whereas the bottom row shows images where defoliation was estimated severely too high (numbers in [%] defoliation, GT = ground truth, PR = predicted, Urmeter = U, Parcours = P, WSI = W): (a) W, GT: 100, PR: 23.4, (b) W, GT: 100, PR: 31.0, (c) W, GT: 100, PR: 45.8, (d) P, GT: 80, PR: 26.5, (e) W, GT: 35, PR: 72.9, (f) P, GT: 30, PR: 65.3, (g) P, GT: 5, PR: 38.7, (h) W, GT: 20, PR: 47.6.

**Figure 9:**
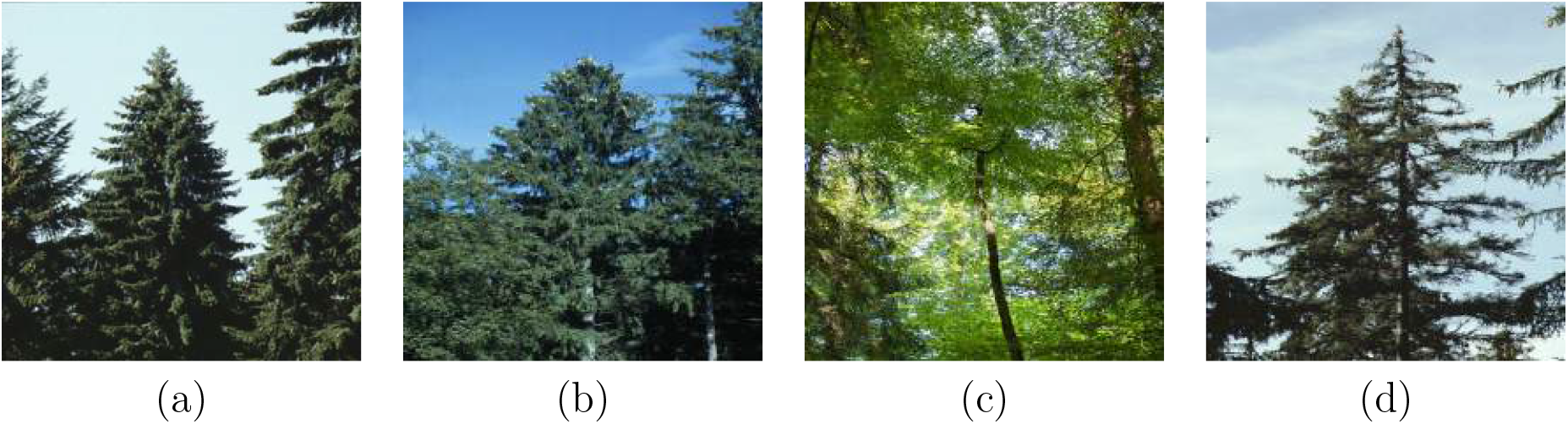
Examples of images where defoliation estimation with our CNN model works very well (prediction errors below 1 percent point): (a) P, GT: 0, PR: 0, (b) U, GT: 10, PR: 10.0, (c) W, GT: 25, PR: 24.5, (d) P, GT: 55, PR: 55.1.

**Figure 10:**
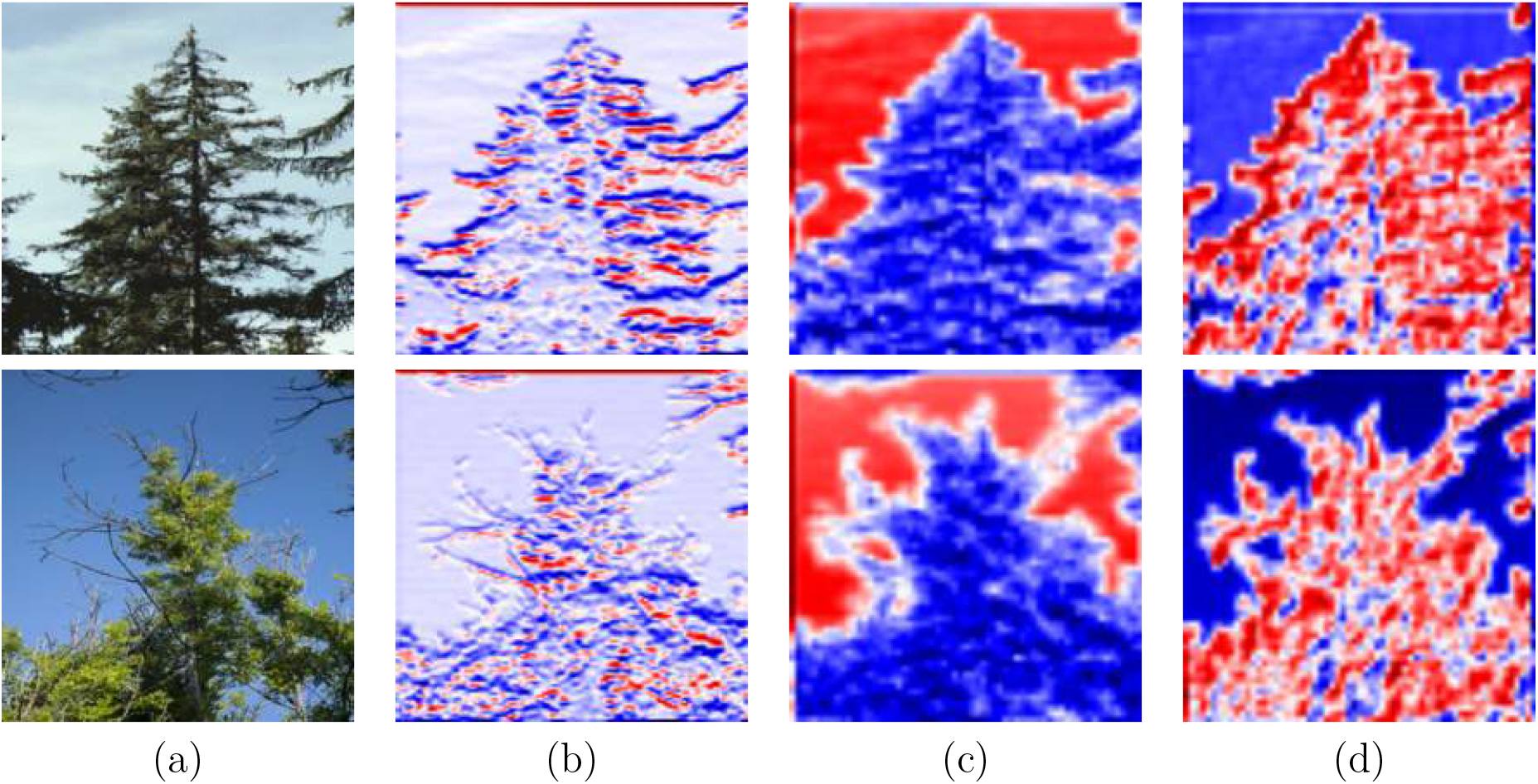
Examples of activation maps: (a) input image (size 256 × 256 pixels), (b) vertical gradient indicating the distribution of branches (after convolution with *conv1* of kernel size 7 × 7 pixels, before batch normalization, activation map size 128 × 128 pixels, one out of 64 activation maps), (c) separation of tree from sky background (both (c) and (d): after last convolutional layer of *conv2*, kernel size 3 × 3 pixels, after batch normalization, activation map size 63 × 63 pixels, two out of 256 activation maps in total), (d) delineation of empty space in the canopy. Dark blue (−1) and deep red color (1) indicate high activations whereas greyish colors indicate low activation.

The top row in Fig. 8 shows cases where the CNN grossly underestimates defoliation. The first three examples in Fig. 8(a)-(c) show trees with 100% defoliation. Both trunks in (a, b) are standing in front of dense, green canopy. Although the CNN does predict quite some defoliation (23.4% and 31.0%) in both cases, it still underestimates significantly, most likely because it erroneously assigns background canopy to the foreground trunk. Fig. 8(c) is challening because the tree of interest does in fact still have leafs albeit only dead (brown) ones, which counts as 100% defoliated. Another hard case is shown in Fig. 8(d), where the defoliated bottom part of the tree crown is mostly hidden by vegetation in the fore-ground. While it seems possible to improve predictions for single, completely defoliated trunks (Fig. 8(a,b)) or trees with only dead leafs that count as 100% defoliated (Fig. 8(c)) if more training data would be available, occluded cases cannot be resolved by the model. A better strategy would be teaching field crews to take photos from viewpoints that allow observing those parts of the tree that are representative for its overall state.

Gross overestimates of defoliation are shown in the bottom row of Fig. 8. Examples (f)-(h) contain conifers with bright backlighting, interpreted by the CNN model as high defoliation. An interesting case is shown in Fig. 8(e), where our CNN grossly overestimates defoliation most likely due to the empty branches sticking out of the canopy. Three activation maps for this image are shown in the bottom row of Fig. 10. Cases with a large number of trees and occlusions like in Fig. 8(h), where a human observer cannot tell which tree is the one of interest, are basically unsolvable by the method. Again, teaching field crews to take photos such that the individual tree of interest can be recognized would be a viable option. Example images of trees where the CNN approach achieves very good performance (Fig. 9) indicate that the model can well handle complex cases, too. We show activation maps of Fig. 9(d) in the top row of Fig. 10. It can be seen that the distribution of branches is extracted via the vertical gradient (Fig. 10(b)), the tree is separated well from the sky in the background (Fig. 10(c)), and empty space in-between branches is captured, too (Fig. 10(d)).

### 4.5. Comparison to related approaches

In this section, we compare our approach and experimental results to related work. Since we are not aware of any other work that estimates tree stress let alone tree defoliation from ground-level images, we discuss pros and cons with respect to research that uses remotely sensed data to assess the state of individual trees.

A general advantage of remote sensing methods is their dense coverage of large-scale regions. Regardless of the sensor technology, airborne LiDAR, aerial or satellite images can provide rich evidence about the health condition of vegetation. While LiDAR can acquire the 3D structure of vegetation with a 3D point cloud directly, remotely sensed multispectral or hyperspectral imagery usually provide evidence through image channels in the infrared domain. While most methods rely on only a single data source like full-waveform LiDAR (Yao et al., 2012a), other methods combine data of different modalities like full-waveform LiDAR and hyperspectral data (Shendryk et al., 2016) or multispectral aerial images with full waveform LiDAR (Polewski et al., 2015a). Yao et al. (2012a) and Polewski et al. (2015a) detect standing dead trees in forests whereas Shendryk et al. (2016) estimate dieback and crown transparency. To distinguish different categories, a Random Forest classifier is used in (Shendryk et al., 2016), a Support Vector Machine in (Yao et al., 2012a), while Polewski et al. (2015a) follow an active learning approach with a logistic regression. All three methods have in common that they base their classification on handcrafted features. They assemble a multi-step workflow with significant human interaction to finally produce a classification output.

Yao et al. (2012a) and Shendryk et al. (2016) have in common that they validate their methods experimentally on very small data sets. Shendryk et al. (2016) train and validate their approach on only 73 individual trees, Yao et al. (2012a) experiment with 0.3 hectares containing 314 trees in total of which 87 are dead trees (under leaf-on conditions). The exact number of trees used for experiments by Polewski et al. (2015a) is not given explicitly. While the authors initially subdivide their 1*km* × 1*km* test plot into 44000 individual trees, they label 3000 trees in three different data sets through visual interpretation of aerial images and finally experiment with 2000 trees including ≈ 436 dead trees. In comparison, we evaluate our method on 2108 tree images with individual defoliation labels acquired in forests all over Switzerland.

Although quantitative results cannot be compared directly due to different data sets, input data, methods and varying degree of human interaction, they can give a rough indication of performance. For their best setup, Yao et al. (2012a) report an overall accuracy of 73%, a kappa score of 0.45, and a misclassification rate of 30%. The best result achieved for dieback in Shendryk et al. (2016) is 81% overall accuracy and a kappa score of 0.66. The highest overall accuracy of 89% is achieved in (Polewski et al., 2015a) (no kappa score is provided) with a multi-step workflow and much manual intervention at different steps of the workflow.

In contrast, our method learns tree defoliation end-to-end directly from the data set without any human intervention. Unlike Yao et al. (2012a), Polewski et al. (2015a), and Shendryk et al. (2016) our workflow is highly automated and does not need any human interaction or even specific, manually fine-tuned data pre-processing at any point. Moreover, our data set comprises 2108 tree images as opposed to (Yao et al., 2012a; Shendryk et al., 2016) who validate their method on only 73 respective 314 individual trees. A continuous defoliation value was assigned to all trees either in situ (*Parcours* and *WSI*) or from the images (*Urmeter*) by experts. Our method regresses a continuous defoliation value, which seems a more challenging task than distinguishing dead trees from healthy ones as done in (Yao et al., 2012a; Polewski et al., 2015a). If training and testing across all three data sets with five-fold cross-validation, we achieve an avgMAE of 7.6% with a very low standard deviation across folds of 0.3%.

In summary, our approach has a higher degree of automation and achieves good results for regressing continuous defoliation values. Our method is quantitatively validated on a bigger data set sampled across a larger area. Nonetheless, it would be desirable to collect further reference data for additional experiments. In comparison, recent works on tree species classification with full waveform LiDAR data uses 3202 (Blomley et al., 2017) respective 9930 (Hovi et al., 2016) individually labeled reference trees. Finally, we use simple RGB images as input data instead of full waveform LiDAR, aerial images or hyperspectral data.

## 5. Conclusions

We have presented a deep machine learning approach for tree defoliation estimation from RGB images acquired with off-the-shelf, hand-held cameras. Adaption of a CNN with a ResNet architecture to tree defoliation shows good results for three out of four data sets of different level of difficulty. As with basically all (deep) machine learning methods, the model performs well for all situations with sufficient training data whereas performance suffers in case the number of training samples is very low. Complex scenes, where highly defoliated trees are photographed in front of dense, green canopy of very low defoliation, can cause errors.

However, despite few, extreme cases where the method fails, the overall performance is very promising. Potential future directions could involve combining ground-level photos with aerial or satellite imagery to densely predict defoliation across large regions. Moreover, WSL is continuing to collect photos during their field campaigns, which will lead to several hundred more images each year. We hope that our approach presented in this paper is one step forward towards making monitoring of forest trees more efficient. Our findings can help providing an additional tool for large-scale forest tree monitoring in the long-term. The approach might allow for assessing many more trees than those few inspected visually by the experts in the field. Taking photos of trees outside the sample plots by the experts can be easily and quickly achieved and multiplies the number of trees with defoliation information in a region. In a next step, crowd sourced images from a citizen science project could be included to increase the spatial coverage of defoliation assessments.

